# Improvement of arthroscopic surgical performance using a new wide-angle arthroscope

**DOI:** 10.1101/399907

**Authors:** Jae-Man Kwak, Erica Kholinne, Kyoung Hwan Koh, In-Ho Jeon

**Affiliations:** Department of Orthopedic Surgery, Asan Medical Center, College of Medicine, Ulsan University, Seoul, South Korea; Department of Orthopedic Surgery, St. Carolus Hospital, Jakarta, Indonesia

**Keywords:** shoulder arthroscopy, wide-angle arthroscope, dimensionless squared jolt

## Abstract

**Background:** We have developed a new arthroscope with a field of view of 150°. This requires less motion to maneuver, and the optical error is decreased. It also improves how novices learn arthroscopy. We hypothesized that surgical performance using this arthroscope is superior to a conventional arthroscope. This study tested the hypothesis using motion analysis and a new validated parameter, “dimensionless squared jolt” (DSJ).

**Methods:** We compared the surgical performance using between the wide-angle and the conventional arthroscope among 14 novice orthopedic residents who performed three standardized tasks three times with each arthroscope. The tasks simulated arthroscopic rotator cuff repair surgical skills. Their motion was analyzed using an optical tracking system. The differences in performance parameters, such as time taken, average acceleration of the hands (m/s^2^), the number of movements, and the total path length (m) including DSJ between the two arthroscopes, were investigated using paired t-tests.

**Results:** All the estimated values for the tasks using the 150° arthroscope were lower than those for the 105° arthroscope. There were statistically significant differences in performance between the two arthroscopes only for DSJ (p = 0.014) and average acceleration (p = 0.039).

**Conclusions:** DSJ and average acceleration are reliable parameters for representing hand-eye coordination. The surgical performance of novice arthroscopists was better with the new wide-angle arthroscope than with the conventional arthroscope.

## INTRODUCTION

Shoulder arthroscopy has evolved and has become an important tool for minimally invasive surgery. As arthroscopic techniques have evolved, arthroscopy has become the gold standard for diagnosis and management of shoulder disorders [1-4]. However, several issues remain, particularly with regards to the long learning curve and demands on the time required for training. Additionally, technical errors arise from the technical errors arising from the limitations of hand-eye coordination required for arthroscopic systems, especially for novice arthroscopists. Several studies have shown that training using a dry or virtual simulator is effective and safe for overcoming these issues [5-7]. However, cost-effectiveness remains a limitation of every training hospital.

Based on our experience, one of the main issues in arthroscopy is the limited field of view (FOV) of conventional arthroscopic systems. Several studies have reported that a limited arthroscopic FOV can make it difficult to identify entire anatomic structures; this was often associated with poor hand-eye coordination regarding surgical triangulation and handling the instrument [8-11]. We hypothesized that a wide-angle arthroscope with a better FOV would result in better hand-eye coordination and would bring important benefits such as an accelerated learning technique, reduction of technical errors, and improvement in the long-term performance of surgical skills.

The aim of this study was to compare hand-eye coordination and the performance of surgical skills between a newly developed wide-angle arthroscope (150° FOV) and the conventional arthroscope (105° FOV). We used motion analysis to evaluate the performance of novice arthroscopists, assessing their hand-eye coordination by using the “dimensionless squared jolt” (DSJ), which is widely accepted as an objective parameter of hand-eye coordination in the field of engineering.

## MATERIALS AND METHODS

### Participants

Fourteen right-handed orthopedic residents, with no prior arthroscopic surgical training, were enrolled in this study.

### Arthroscopic systems

A conventional 30° arthroscopic system (IM4000, IM4120; ConMed Linvatec, Utica, NY, USA) was used to perform the tasks in this study and compared with a new wide-angle arthroscopic system (MGB Endoscopy Company, Seoul, Korea) that had a high-definition lens with a 150° Field of View (FOV).

### Arthroscopic tasks

The participants performed a set of tasks using the two arthroscopes sequentially in a randomly allocated order. A Latin square counterbalancing technique was used to compensate for learning effects from previous practice [12]. Prior to the experiment, each participant was given an instructional video manual that covered all the experimental processes and arthroscopic tasks to be performed, as well as an introduction to the experimental and surgical instruments. The participants were allowed 10 minutes to become familiar with the experimental system and surgical instruments. The participants performed the tasks without any assistance.

A dry right shoulder model (Arthrex, Naples, FL, USA) was used for the tasks (**Fig 2a**). Black nylon sutures were made at five sites along the lateral border of the rotator cuff, and the bicipital groove was marked in blue. A pilot hole for an anchor was made at an appropriate position on the greater tuberosity (**Fig 2b**). The model and related instruments were set up on a table, with a pair of reflective hand illustrations placed on the table to mark the start and end positions for the participants (**Fig 4b**).

**Fig. 1.**
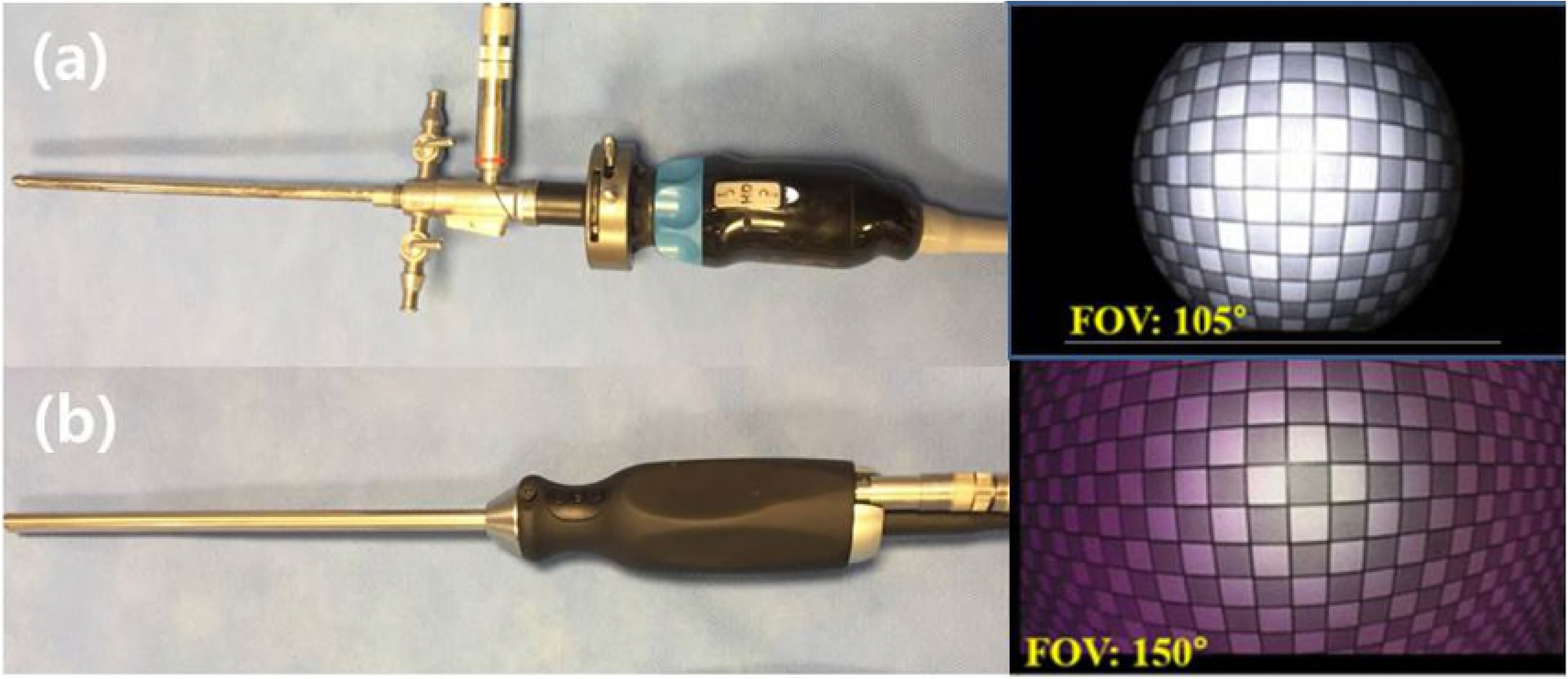
The two arthroscopes used in this study, with representative images showing the field of view. **(a)** The conventional arthroscopic system. **(b)** A new wide-angle arthroscopic system.

**Fig. 2.**
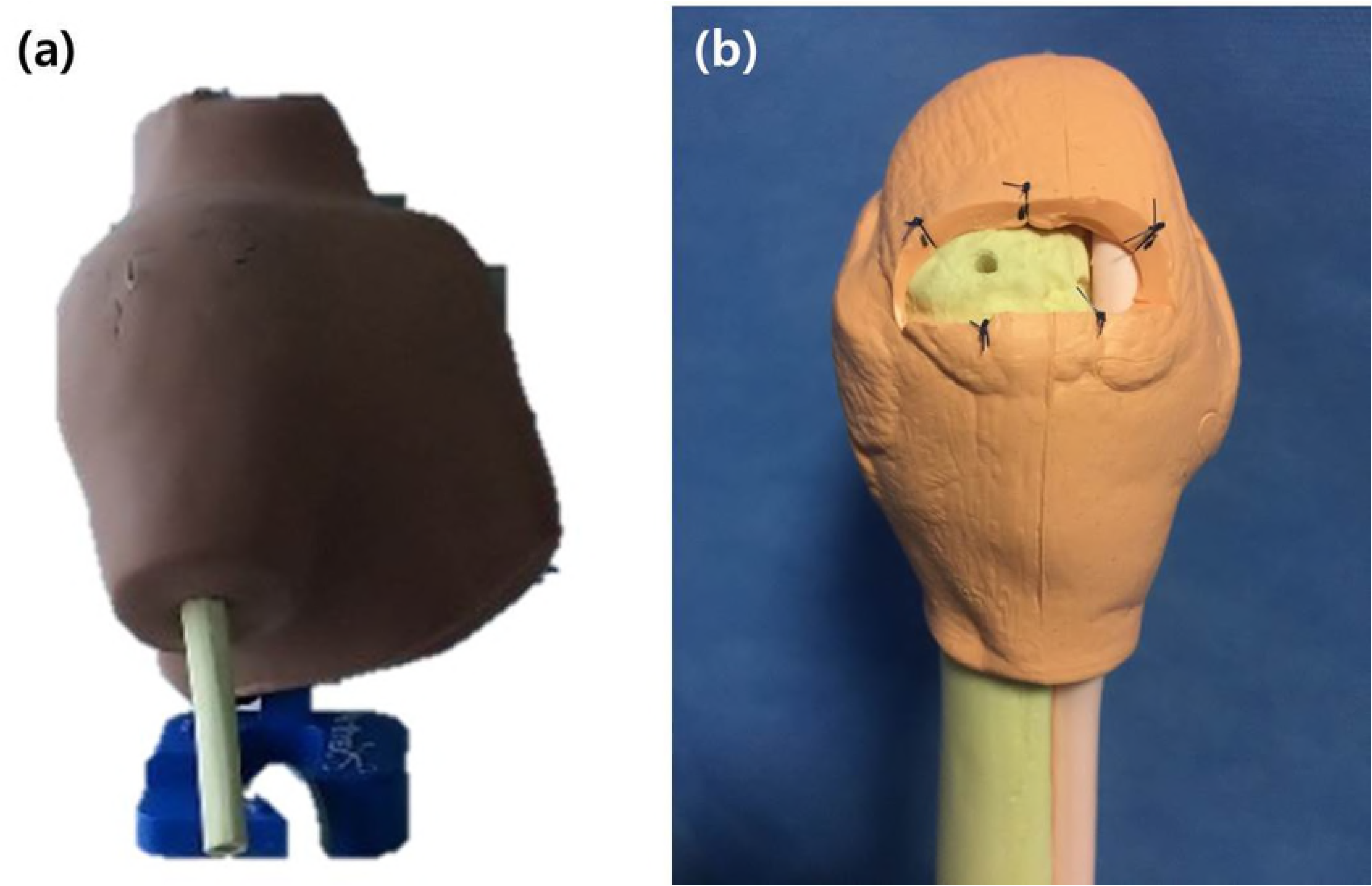
**(a)** A dry human shoulder model was used for the arthroscopic tasks. **(b)** Black nylon sutures were made at five sites along the lateral border of the rotator cuff (*red arrows*), and the bicipital groove was marked in blue. A pilot hole for an anchor was made at an appropriate position on the greater tuberosity.

The three assigned tasks, which simulated basic surgical techniques, were as follows: (a) touching the five points along the rotator cuff marked by sutures twice using the grasper, passing through the anterior portal; (b) inserting an anchor at the predetermined point on the footprint of the rotator cuff; and (c) pulling the suture through the anterior portal using the grasper (**Fig 3a-3c**). These three tasks have been validated by previous studies [6, 13-15]. Each task was performed three times with each scope.

**Fig. 3.**
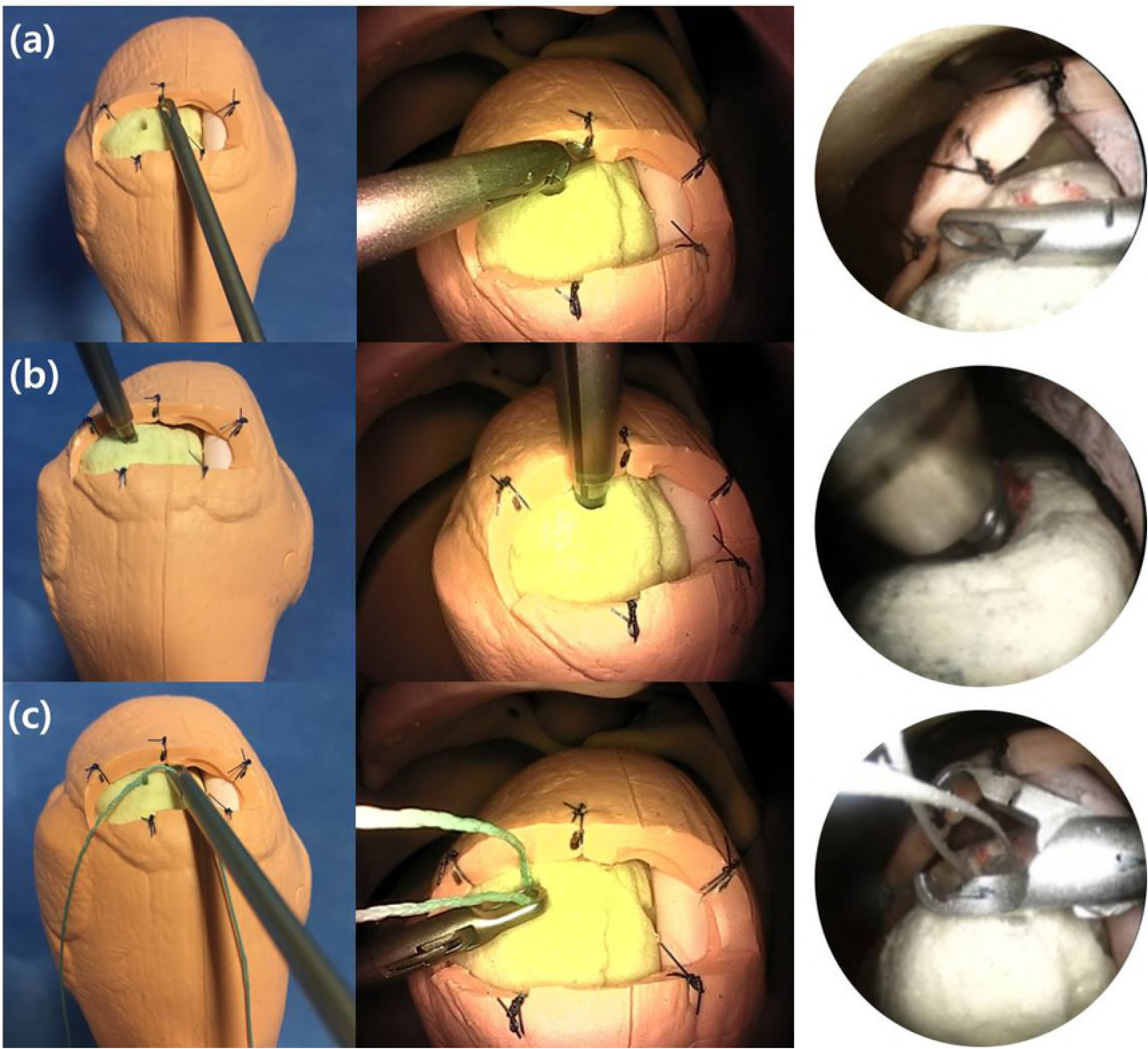
The three arthroscopic tasks. **(a)** Touching five points on the rotator cuff twice using the grasper, passing through the anterior portal. **(b)** Anchor insertion at a predetermined point on the footprint of the rotator cuff. **(c)** Pulling the suture through the anterior portal using the grasper.

### Hand motion analysis

During the tasks, the participant’s hand movement was tracked by an optical motion analysis system comprised of eight motion-capture cameras (Prime 41; Natural Point, Inc., Corvallis, OR, USA) (**Fig 4a**), 10-mm reflective marker balls placed on the third metacarpal area of the dorsal side of participant’s hands, and tracking software (Motive: Tracker; Natural Point, Inc.) (**Fig 4b**). The system continuously recorded three-dimensional (3D) data for the two reflective markers on the participant’s hands. The following data were collected and analyzed using motion analysis, as described in previous reports [10, 14, 16]: (1) the time taken (in seconds) to complete the tasks, recorded from the time the participant started to lift their hand from the table until the time the task was completed and the participant’s hand was resting on the table; (2) the average acceleration of the hands (m/s^2^); (3) the number of movements made, defined by changes in velocity with respect to time that exceeded the predetermined threshold value of 10 m/s^2^; and (4) the total path length (m), the total distance traveled by the reflective marker during the task.

**Fig. 4.**
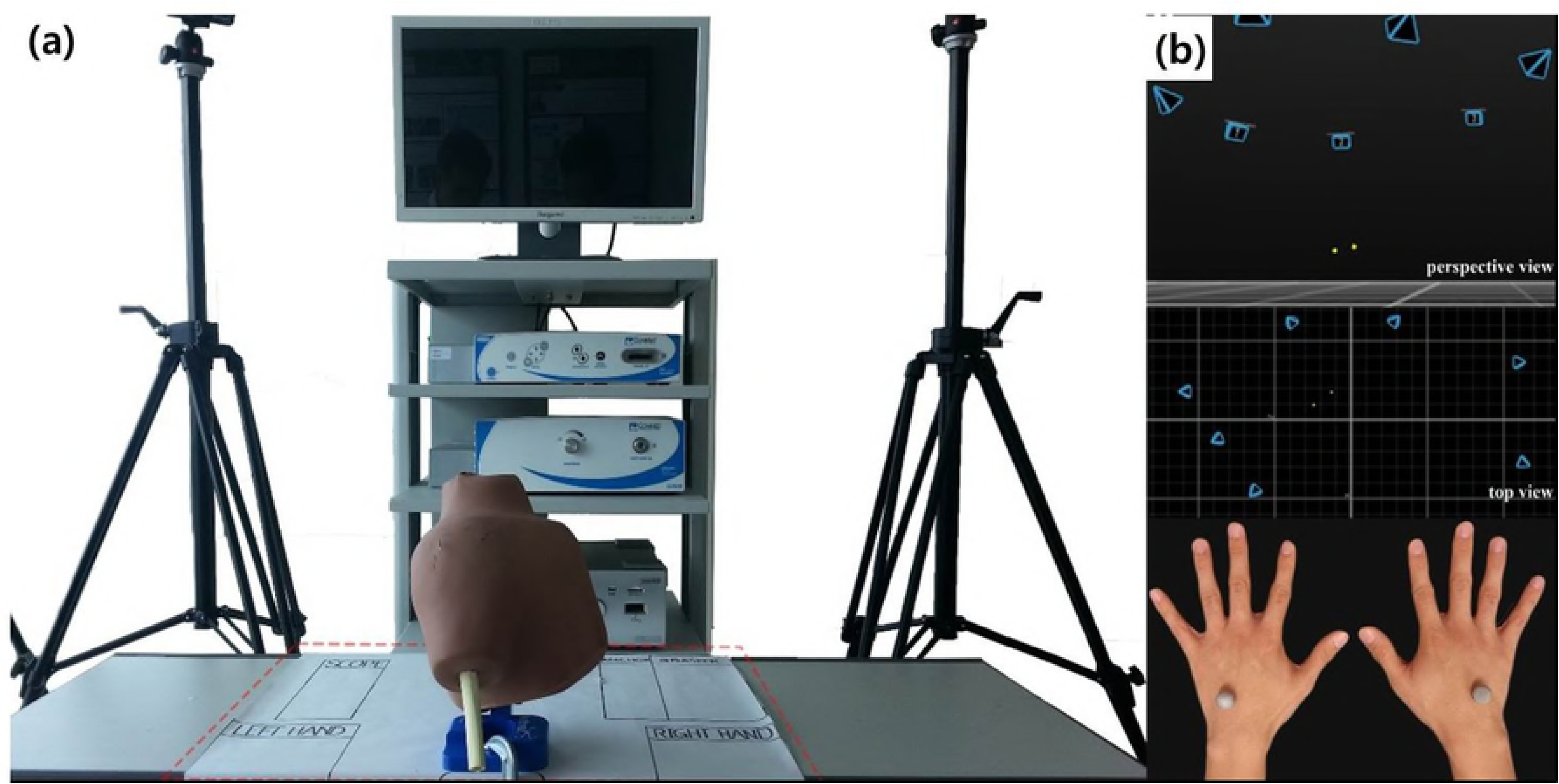
The optical motion analysis system, which was comprised of eight motion-capture cameras, reflective markers placed on the third metacarpal area of the dorsal side of the participant’s hand to track hand motion, and tracking software.

Using these data, DSJ, a physical index for estimating hand-eye coordination, was calculated by MATLAB 2012b (Math Works, Torrance, CA, USA) using the following formula:

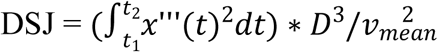

where D is the duration of the movement and *v*_*mean*_ is the movement’s average velocity.

### Statistical analysis

The time taken to complete the tasks, the total path length, the number of movements and the average acceleration data, including DSJ, were normally distributed and categorized as continuous variables. However, DSJ and average acceleration were not normally distributed. An *a priori* power analysis showed that a sample size of 34 attempts using each arthroscope would have sufficient power to show a significant difference, assuming a significance level of 0.05 and power of 80%. The *p*-values for time to complete, total path length, and the number of movements were calculated using a paired t-test, with the Wilcoxon signed rank test used for DSJ and average acceleration; *p* < 0.05 was considered to be statistically significant. The analyses were performed using the SPSS software package (version 20.0; IBM Corp., Armonk, NY, USA).

## RESULTS

DSJ (p=0.014) and average acceleration (p=0.039) were the only parameters which showed a statistically significant difference between the two arthroscopes. There were no other statistically significant differences between the arthroscopes, including the time taken (p=0.282), the number of movements (p=0.323), or total path length (p=0.142). All other estimated values for the tasks using the 150° arthroscope were lower than those for the 105° arthroscope. The collected data are summarized in **Table 1** and **Figure 5.**

**Fig. 5.**
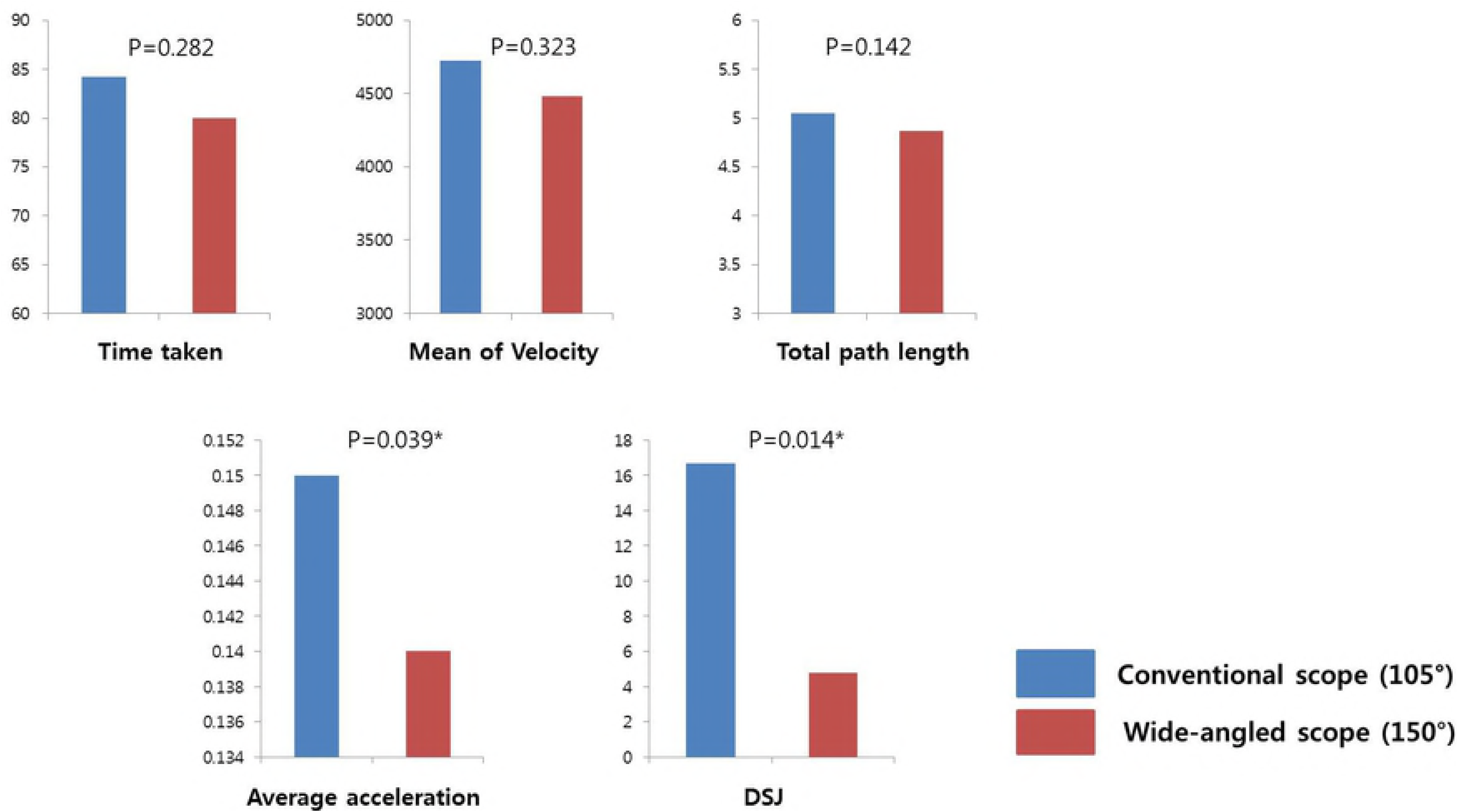
Bar charts comparing the estimated values of parameters between the conventional 105° FOV arthroscope (*blue*) and the 150° FOV arthroscope (*red*).

## DISCUSSION

Previous studies have considered and demonstrated that improving the arthroscopist’s view results in better performance; in other words, improved hand-eye coordination leads to better surgical performance [8-10, 16-21]. Based on this theory, we introduced the prototype of the wide-angle arthroscope in a previous study [9]. Compared to conventional arthroscopes, which have a 105° FOV, the new arthroscope with a 150° FOV provides unprecedented wide-angle viewing. However, our hypothesis demonstrated scientifically that this would not simply result in superior performance because it involves various complicated movements; the main issue is identifying a specific tool to estimate movement quality. The global rating scale, which has been widely used, is an efficient and easy way to estimate surgical performance [22-25]. However, assessing subjective factors remain because arthroscopy is conducted under the observation of another person [26].

To overcome this subjective limitation, we tried to objectively find the tool to assess hand-eye coordination, which is highly related to the surgical performance. Fortunately, motion tracking systems make it possible to obtain objective data regarding hand movements [14, 27-30]. Using the motion tracking system, we evaluated the participants’ performance and hand movements by the time taken to complete the tasks, the total path length traveled by their hands, the number of movements made, and the average acceleration of the hands. From these, we calculated DSJ, a relatively simple indication of movement quality. A “jolt,” also known as a “jerk,” is a physical value indicating the rate of change of acceleration for a movement. In DSJ, this is squared to counterbalance any negative values and is presented as a dimensionless parameter to allow it to be comparable. DSJ has previously been used to evaluate several movement disorders [31, 32] and has been regarded as an effective parameter for quantifying the quality of movement [33, 34]. Hogan *et al*. [32] noted that “a dimensionless jerk-based measure properly quantifies common deviations from smooth, coordinated movement.” This suggests that appropriate measurement of DSJ would reflect the quality of hand movements of the arthroscopists.

From the results of our previous study which demonstrated the validity of DSJ for objective assessment of hand-eye coordination, we can use the parameter to objectively assess surgical performance.[35] In the present study, DSJ and average acceleration showed statistically significant differences between the results for the two arthroscopes. Based on the results, we finally demonstrated objectively the superiority of the new wide-angle arthroscope compared with the conventional arthroscope.

One limitation of this study included the relatively small sample size. In addition, the arthroscopic surgical tasks were performed using a dry model, which does not perfectly reflect the clinical situation. Nevertheless, we believe that the new wide-angle arthroscope may improve arthroscopic performance and offer a superior training tool compared with the conventional arthroscope.

## CONCLUSION

The surgical performance of novice arthroscopists was improved with the wide-angle arthroscope as compared with the conventional arthroscope.

### Disclosure

This research was supported by a grant of the Korea Health Technology R&D Project through the Korea Health Industry Development Institute, funded by the Ministry of Health & Welfare, Republic of Korea (grant number: HI13C1634). The funding source had no involvement in the study design; collection, analysis, and interpretation of data; writing of the report; or decision to submit the article for publication.

